# Self-organized criticality in a mammalian RNA-interference circuit

**DOI:** 10.64898/2026.11.09.765295

**Authors:** Jordi Pla-Mauri, Marc Güell, Ricard Solé, Núria Conde-Pueyo

## Abstract

Living systems have been argued to operate near critical points, but their relevance at the cellular level is poorly understood. Critical states allow for a high sensitivity to external inputs and maximal information transfer and display fluctuations at all scales. Critical points are very special, and require the fine tuning of a *control parameter*, which is hard to reconcile with the robustness of living cells. Self-organized criticality (SOC) proposes that feedback, rather than tuning, brings a system to its critical point. Here we show that RNA interference, in which messenger RNAs compete for shared silencing machinery and so queue for service, together with negative feedback gives rise to SOC in mammalian cells. In a synthetic circuit, a protein repressor whose mRNA competes with the mRNA of a fluorescent reporter that it represses senses the congestion of the queue, closing the feedback loop. Theory and stochastic simulations show that the loop holds the queue near its critical point despite changes in its parameters, and that expression becomes intermittent, with scale-free bursts. Measurements in human cell lines show the predicted power-law distributions of single-cell fluorescence. RNA interference could thus support engineered SOC in animal cells and may also contribute to SOC in natural systems.

## I. INTRODUCTION

Physical systems close to a critical point respond to small perturbations with large, collective changes, and exhibit fluctuations without a characteristic scale [1–3]. Familiar critical points, such as the liquid-gas critical point or the Curie point of a magnet, require tuning a control parameter to a particular value. In 1987, Bak, Tang and Wiesenfeld proposed that some systems might spontaneously approach criticality without such external tuning [4, 5]. In sand or rice piles, grains are slowly added and redistributed through avalanches. The driving force increases the slope, while avalanches relax it, potentially maintaining a marginal state characterized by events spanning many scales [6, 7]. This phenomenon, known as *self-organized criticality* (SOC), was subsequently invoked to explain the dynamics of forest fires [8], ecosystems [9, 10], and evolution [11, 12].

Yet, establishing whether natural systems genuinely self-organize toward criticality remains a major challenge. Their underlying interactions, environmental drivers, and feedback mechanisms are often difficult to disentangle, making it hard to distinguish causal mechanisms from statistical signatures. Synthetic biology offers a complementary route to this problem: rather than inferring the existence of SOC from observations of complex natural systems, we can reconstruct its essential ingredients in experimentally controllable biological systems. Theoretical frameworks suggest that SOC can emerge through feedback between an order parameter, which characterizes the collective state of a system, and a control parameter that is dynamically driven toward a phase transition [13–19]. This perspective suggests a concrete experimental strategy: identify a biological system exhibiting a genuine phase transition, engineer a molecular sensor of its collective state, and couple that sensor to a feedback circuit controlling the distance to criticality. By constructing, perturbing, and rewiring these components, synthetic biology can transform SOC from a hypothesis inferred from natural patterns into a mechanistic principle that can be directly tested.

Critical states are of particular interest in living matter. A system at a critical point has maximal sensitivity, the widest dynamic range and long-range correlations, properties that are valuable for sensing, computation, and adaptation [20–23]. Evidence has been reported at very different levels of organization: avalanches of activity in cortical tissue [24–27], scale-free correlations in bird flocks [28, 29], Internet fluctuations [30, 31], traffic jams [32, 33] and gene regulatory networks that sit near the border between order and chaos [34–36], with local adaptive rules [37] and selection for evolvability or information [38, 39] offered as routes by which such states could be reached (for reviews see Refs. [40, 41]).

However, evidence for self-organized criticality (SOC) has often relied primarily on the observation of power-law distributions. Such evidence is insufficient: heavy-tailed distributions can emerge from mixtures of states, multiplicative noise, or observational thresholds, without the system being near a critical point [42–45]. Establishing SOC therefore requires identifying the mechanisms that drive and maintain a system near criticality. Ideally, these mechanisms should be experimentally accessible, allowing the transition, sensing, and feedback processes to be independently manipulated. A molecular system whose components can be dosed, deleted, or rewired would provide a particularly powerful test of SOC as a causal mechanism rather than a statistical signature.

Gene expression is a useful setting for such a system. Its machinery is discrete, stochastic and shared; the resulting fluctuations generate phenotypic diversity, bet hedging and fate decisions [46–51]. Using synthetic biology, it has been possible to build circuits from a few characterized parts [52, 53] and shown that limited pools of shared resources, such as polymerases, ribosomes and proteases, couple otherwise independent genes [54–57]. Engineered controllers can compensate for this coupling [58, 59]. Shared degradation is a clear example. When two substrates compete for one saturable enzyme, they queue, and one substrate’s queue delays the removal of the other [60]. This queue has a phase transition: below the enzyme’s capacity it stays short and flows freely; above the capacity, it grows; at the balance point, its length fluctuates at all scales. Vidiella and colleagues used this principle with the ClpXP protease of *Escherichia coli*. A negative feedback that senses protease congestion turned a two-gene circuit into a self-organized critical system with heavy-tailed, bursty expression [61]. Whether the same mechanism works in mammalian cells, with their different machinery, noise and timescales, is unknown.

Mammalian cells have their own shared and saturable machinery for removing messenger RNA: RNA interference. A short hairpin RNA (shRNA) is processed into a guide that loads Argonaute2; the resulting RNA-induced silencing complex (RISC) recognizes a complementary transcript, cleaves it and is released to act again [62–67]. Because the enzyme turns over at a finite rate, all transcripts that carry the target compete for it. This competition is measured as the dilution of silencing by target abundance [68] and by transfected small RNAs [69], as thresholds in target expression [70], as lethal over-saturation of the pathway [71], and as crosstalk between endogenous transcripts that share microRNAs [72–76]. RNA interference therefore acts as a queue, with messages as customers and RISC as the server. It has already been combined with repressor proteins in tunable switches [77–79], but the queue has not been used to set its own operating point.

In this work, we implement this design on a single plasmid. The messages for LacI and a fluorescent reporter carry the same shRNA target and share one queue. As the queue lengthens, the LacI message lives longer, so LacI protein senses the waiting time. LacI represses the reporter promoter, whose message also joins the queue. aTc controls the machinery capacity and IPTG sets the sensor’s operating point. We show that the queue has a critical point, the loop holds the circuit near it as drive, capacity, and dose change, and the resulting operating point produces avalanche-like excursions with power-law amplitudes. Stable HeLa lines carrying EGFP and d2EGFP reporters show tails over more than a decade, with exponents that vary little between reporters.

## II. RESULTS

### A. A saturable RNAi pathway is a queue with a critical point

The circuit is shown in Fig. 1(a). A single plasmid carries four transcription units: the LacI repressor, an shRNA whose promoter is repressed by TetR, TetR itself, and a fluorescent reporter, EGFP or its short-lived variant d2EGFP, under a promoter that LacI represses. The *lacI* and reporter messages carry the shRNA target in their 3^′^ untranslated regions and are therefore cleared by the same RISC. Two inducers control the circuit: aTc releases the TetR shRNA promoter and so increases the amount of RISC, that is, the silencing capacity, and IPTG releases LacI from its operators.

**FIG. 1.**
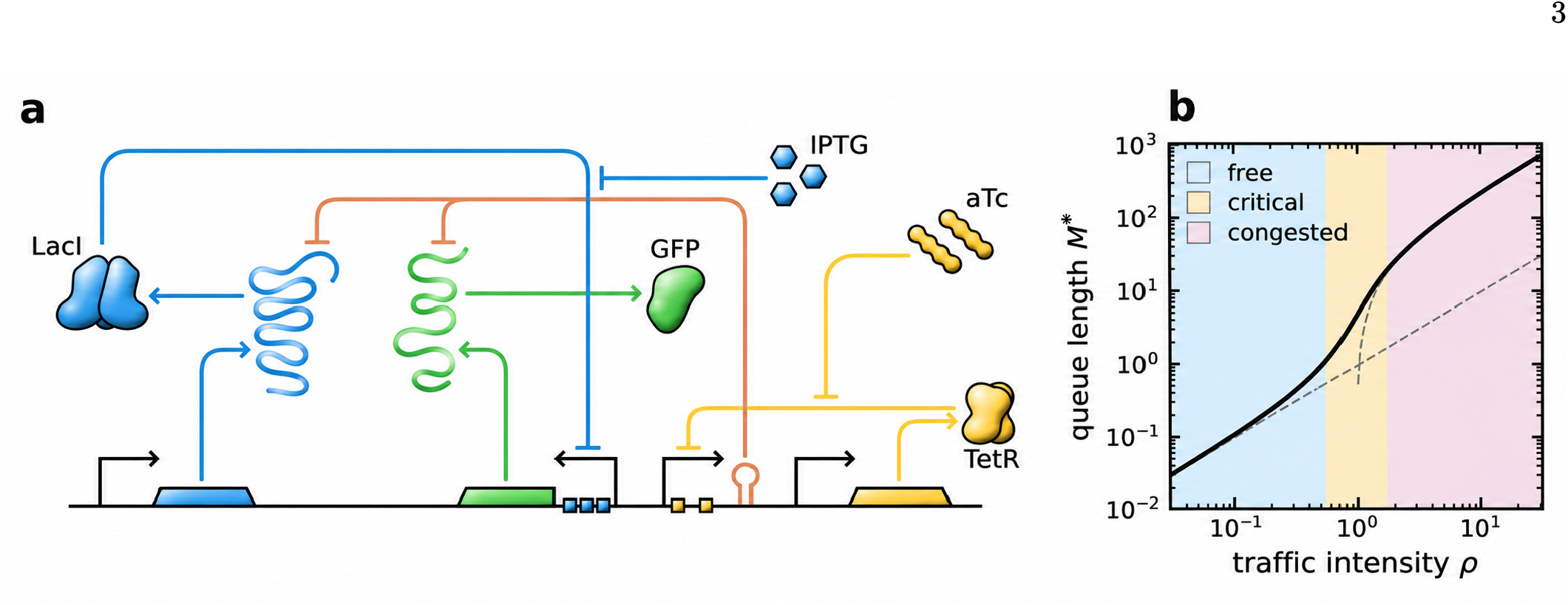
A single-plasmid circuit that queues for RNA interference. (a) LacI (blue) represses the reporter promoter, which carries *lacO* operators; IPTG inhibits LacI. A U6-tetO cassette expresses an shRNA (orange) that silences both the *lacI* and the reporter message through a shared 3^′^ target; TetR (yellow) represses the cassette and aTc inhibits TetR. The reporter is EGFP or d2EGFP. Both messages are substrates of the same silencing complex, and so they queue for it. (b) Expected queue length *M* ^*^, in units of the Michaelis constant of the silencing complex, against the traffic intensity *ρ*, the demand for silencing divided by its capacity (solid line, reference capacity). The dashed lines are the two limiting behaviors: a short queue proportional to demand when demand is far below capacity (*M* ^*^ ≃ *ρ*), and a queue that grows with the excess of demand when it is far above (*M* ^*^ ≃ *η*(*ρ* ™ 1)). The critical band, where the queue responds most strongly to demand, is the crossover between them.

RISC is modeled as a Michaelis-Menten enzyme shared by both messages (Supplementary Material (SM), Sec. S1). Time is measured in dilution lifetimes and message concentrations in units of the Michaelis constant. If *M* is the total amount of target-bearing message, each message is lost at its basal rate multiplied by

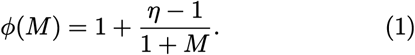

The parameter *η* = *ϕ*(0) is the capacity of the machinery, that is, the fold knockdown of a target in an empty queue, as reported by a standard shRNA knockdown assay. As messages accumulate, the enzyme saturates and each message lives longer, *ϕ* → 1. If both messages are produced at a total demand *A*, the mean queue length that balances demand and clearance, *A* = *Mϕ*(*M* ), is

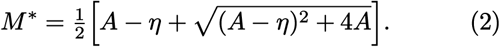

Its behavior is governed by the traffic intensity, the ratio of demand to supply,

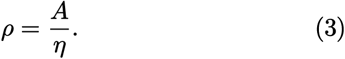

For *ρ* ≪ 1 messages are cleared on arrival, so the queue is short and proportional to demand; for *ρ* ≫ 1 the enzyme is saturated and excess demand accumulates,

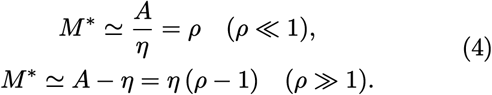

These are the free and the congested phases [Fig. 1(b)]. They meet at *ρ* = 1 in a rounded crossover, which sharpens into a kink as the capacity grows. The queue responds to demand through a change in the mean time that a message spends in the system, the waiting time (Little’s law [80]),

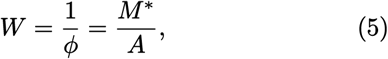

in units of the basal message lifetime. The relative size of the response,

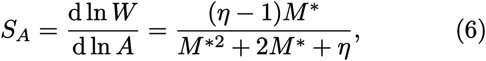

is the susceptibility of the queue, which measures the sensitivity of the waiting time to the arrival rate. It is largest exactly at *ρ* = 1, where it grows without bound with capacity, so the critical point of the queue is the point where demand equals supply. We define the critical band as the interval in which *S*_*A*_ exceeds half its maximum. It is centered on *ρ* = 1 and narrows as the transition sharpens:

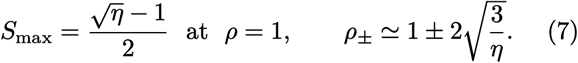

Figure 2(a) shows where the three regimes lie in the plane of demand and capacity: the critical band is a narrow wedge around the line *A* = *η*.

**FIG. 2.**
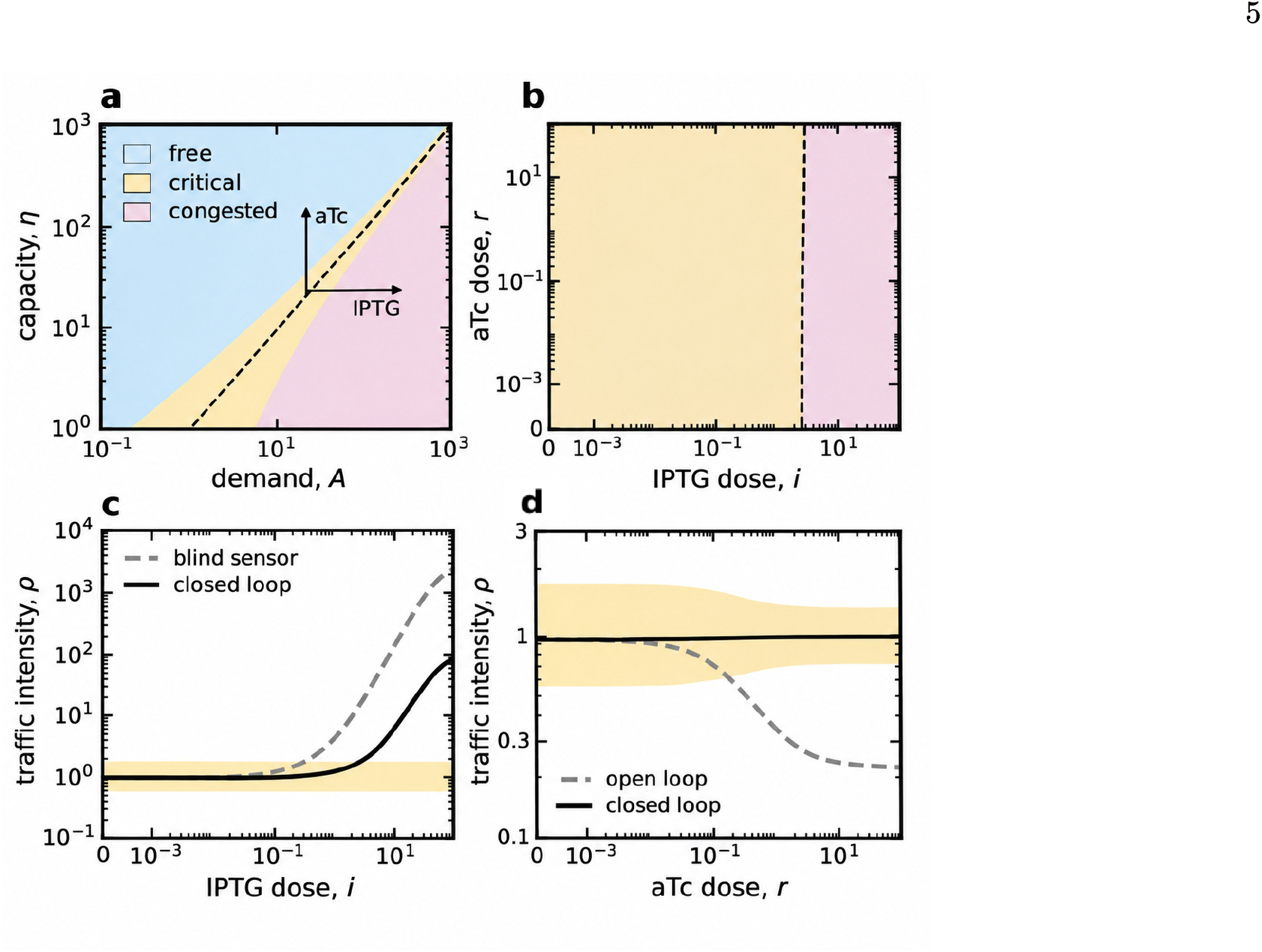
The queue has a critical point and the loop holds the circuit at it. (a) Queue regimes in the demand *A*-capacity *η* plane; the dashed diagonal is *A* = *η*. The arrows show how IPTG raises demand and aTc raises capacity. (b) Closed-loop regimes across IPTG and aTc doses. No free-phase region appears: at zero IPTG the reference circuit is already in the critical band, and more IPTG weakens LacI repression and raises demand toward congestion; aTc changes capacity but, for the circuit’s *n*_*I*_ = 2, leaves the closed-loop operating point nearly independent of capacity. The dashed contour marks the upper edge of the critical band. (c) Traffic intensity against IPTG dose. The blind sensor is repressed by LacI like the closed loop but its LacI message does not read the queue; its reporter drive is set to match the closed loop’s deterministic demand at zero IPTG. It cannot adjust its demand as the queue changes. (d) Traffic intensity against aTc dose, which raises capacity from its basal value *η*_0_ to maximum *η*_∞_. The open-loop control has a fixed promoter that ignores LacI, calibrated to match the closed-loop deterministic demand at zero aTc. In (c,d), black solid is closed loop, gray dashed is the control, and shading marks the critical band. Dose axes are linear below 10^™3^ and logarithmic above. All panels show deterministic fixed points with reference parameters (SM Sec. S1).

#### Open and closed loops

The narrowness of this wedge is the fine-tuning problem of ordinary criticality. In an *open loop*, the reporter promoter is not controlled by LacI: its demand is preset and does not respond to changes in the length of the queue. Such a circuit is critical only if the fixed demand happens to match the capacity, and it leaves the critical band whenever either changes: a stronger promoter pushes it toward congestion and more silencing capacity toward the free phase (arrows in Fig. 2a). In a *closed loop*, LacI reads the queue through its message lifetime and changes the reporter demand in response. Each closed-loop circuit is compared with an otherwise identical open-loop control whose promoter is set to the same deterministic demand at the reference condition. This comparison tests how feedback changes the response to subsequent perturbations; it does not require the stochastic reporter distributions to have the same mean.

### B. Waiting-time sensing drives the queue to its critical point

LacI is translated from a message that is itself cleared by the queue. LacI protein, which is lost only by dilution, integrates the messages over a dilution lifetime, so at fixed demand its concentration is

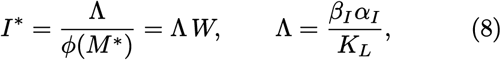

where *α*_*I*_ is the demand of the *lacI* message, *β*_*I*_ the LacI translation rate, and Λ the LacI level of an empty queue in units of the concentration *K*_*L*_ that half-represses the promoter. LacI is thus a proportional read-out of the waiting time in the queue. It represses the reporter promoter with cooperativity *n*_*I*_ = 2, as a LacI tetramer with two DNA-binding heads closes a loop between two operators [81, 82]. The promoter activity is

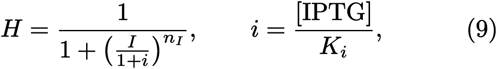

where *i* lowers the active LacI fraction. The loop is closed: more load, longer LacI message lifetime, more LacI, less reporter demand, less load.

The loop imposes a set point. At the fixed point the promoter is repressed by exactly the depth the queue can clear,

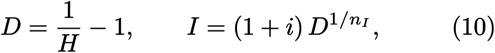

so Eq. (8) fixes the clearance rate the loop holds,

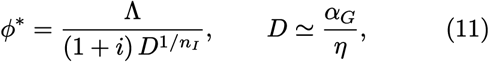

with *α*_*G*_ the strength of the reporter promoter. A held clearance rate corresponds to a definite traffic intensity,

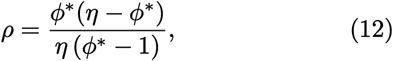

which is exactly one at 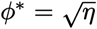 and stays between two thirds and three halves for every clearance rate from 3 up to a third of the capacity. Writing 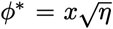, Eq. (12) becomes a function of *x* alone for large *η*, and Eq. (11) identifies *x* with the combination

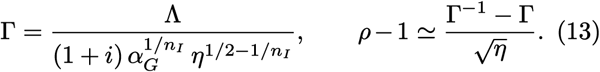

The operating point of the whole circuit is set by this single combination of parameters. The queue is critical at Γ = 1 and stays inside the critical band for 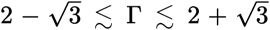 in the large-capacity approximation; larger Γ is the free phase and smaller the congested one. The zero-inducer reference value follows from model parameters calibrated against the EGFP tail; no inducer was added to set the experimental operating point. The combination depends on promoter strength only as 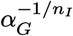 and, for the cooperativity *n*_*I*_ = 2 of the circuit, not at all on capacity:

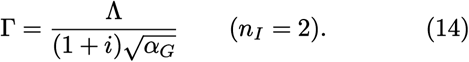

The loop therefore keeps the queue critical while the promoter strength varies over the factor 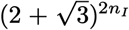, the LacI excess Λ (or the factor 1 + *i* by which IPTG divides it) varies over 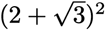, and aTc changes the capacity, provided that *η* ≫ 1. A reporter that ignores LacI has instead

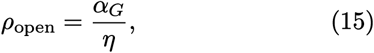

and stays in the band only for a narrow window of either. The feedback compresses a change of promoter strength by the loop gain, which is largest inside the critical band because the susceptibility is (SM Sec. S2).

The deterministic solution of the full circuit confirms these relations (Fig. 2). When IPTG weakens LacI repression, the closed-loop traffic intensity changes far less than that of a sensor blind to the queue and stays in the critical band over the window given above, whereas the blind sensor leaves it at a small fraction of that dose [Fig. 2(c)]. When aTc raises capacity several-fold, the open loop falls into the free phase while the closed loop barely moves, because Γ does not depend on capacity [Fig. 2(d)]. IPTG divides Γ by 1 + *i*, so the closed loop stays critical across aTc doses until 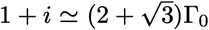, where Γ_0_ is its value at zero inducers; beyond that LacI can no longer close the promoter and the circuit becomes congested [Fig. 2(b)]. The feedback therefore confines the circuit to a band that extends across the IPTG-aTc plane, from the critical regime into congestion but not into the free phase.

### C. Sensor noise turns the pinned state into avalanches

The fixed point is a mean, not a trajectory. Messages and proteins are present in few copies, so that the loop operates through discrete random events. We simulated the message and LacI reactions exactly with Gillespie’s algorithm [83], and calculated reporter protein as the exact conditional mean filter of its message trajectory (Methods and SM Sec. S1). We compared four conditions of the same circuit (Fig. 3). In the *free* phase (Γ ≫ 1) the reporter promoter is too weak to load the queue, which stays nearly empty; the reporter, made from a few messages, fluctuates moderately. In the *congested* phase (Γ ≪ 1) high IPTG keeps the promoter open, the queue is long, and the reporter, high and slow, fluctuates little relative to its level. The *critical* circuit (Γ ≃ 1) is different: its traffic intensity keeps returning to the critical band and makes excursions far above it, and the reporter is intermittent, with long quiet periods and bursts spanning more than an order of magnitude. The mechanism is visible in the traces. When LacI messages are transiently absent, LacI protein decays at the dilution rate, repression relaxes and the promoter opens; the queue congests and the messages live longer; the next LacI message is then translated into a larger burst that closes the promoter. Slow drive, threshold and fast relaxation are the ingredients of an avalanche. Finally, the *open loop*, matched to the same deterministic demand at the reference condition, is critical only at that condition: its reporter fluctuates weakly around a constant, and its traffic intensity does not move, because nothing feeds the state of the queue back to the demand.

**FIG. 3.**
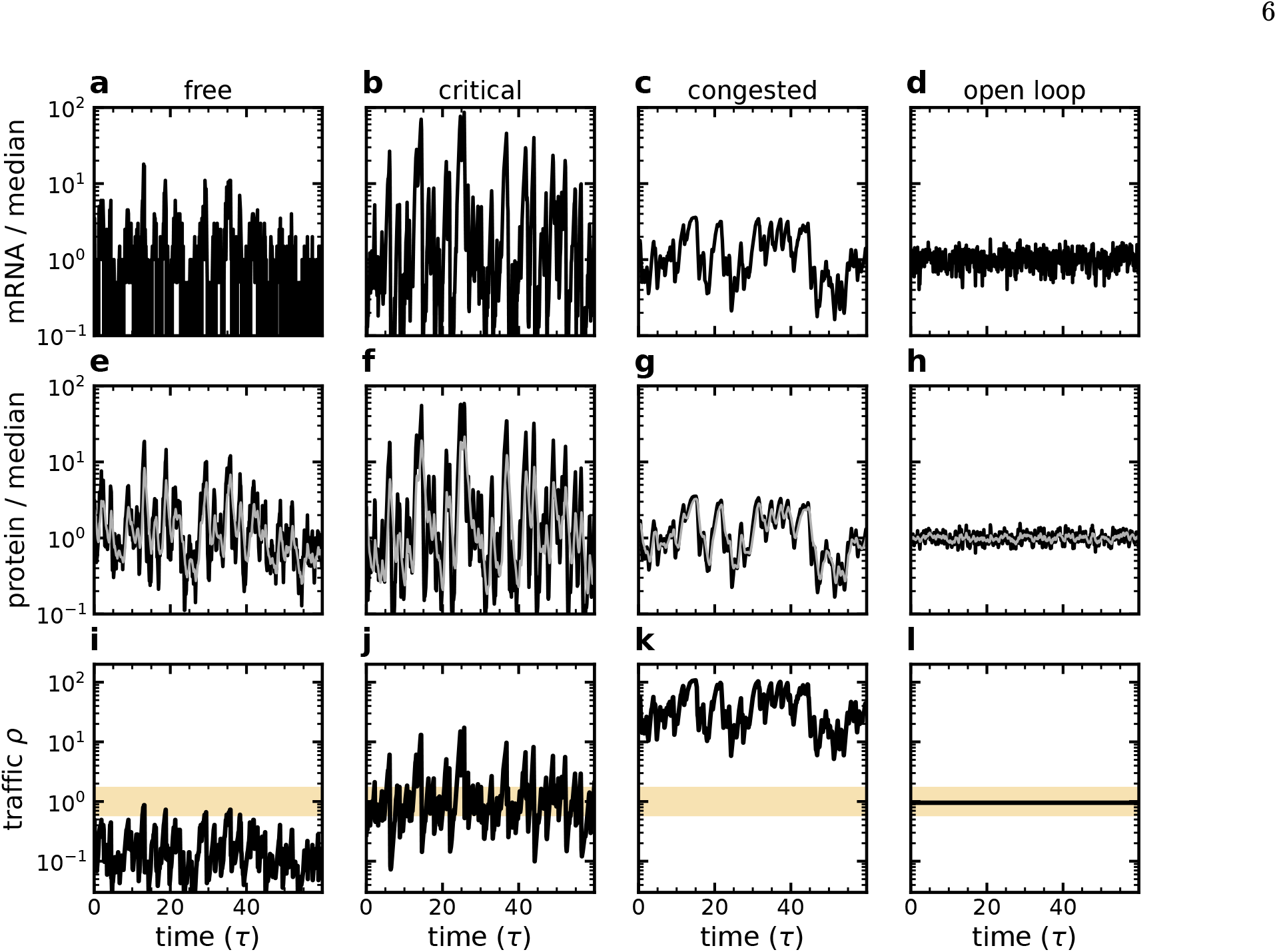
Avalanches appear when the loop holds the queue at its critical point. Top row, reporter mRNA divided by its median. Middle row, EGFP (gray) and d2EGFP (black) protein, each divided by its own median; reporter protein is the conditional mean filter of the shared mRNA trajectory. Bottom row, instantaneous traffic intensity *ρ*(*t*) with the critical band shaded. Time *τ* is in dilution lifetimes. Columns show the free phase (weak reporter promoter, Γ ≫ 1), the critical reference circuit (Γ ≃ 1), the congested phase (high IPTG, Γ ≪ 1), and an open-loop control whose reporter promoter ignores LacI and is set to match the critical case’s deterministic demand at the reference condition; its traffic intensity is constant by construction. Each column uses one Gillespie trajectory for the messages and LacI protein; free and congested parameters are in SM Sec. S1.

The amplitudes of the excursions are scale free over a finite range (SM Sec. S3), and the model gives a mechanism for the exponent. A stable protein made in bursts at frequency *κ* per dilution lifetime decays as *e*^™Δ^ during a burst-free interval Δ, which has probability *e*^™*κ*Δ^; hence *P* (*I < x*) ∝ *x*^*κ*^ at low *I*. The reporter message follows LacI with a local log-slope

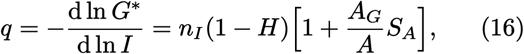

where *A*_*G*_ is the part of the demand that comes from the reporter. If the low-*I* sensor regime and an approximately constant-*q* transfer window overlap, that sensor power law is carried into the reporter with

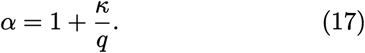

Here *κ* is the frequency at which the *lacI* gene fires. The factor *S*_*A*_ is the contribution of the queue, which is largest at the set point:

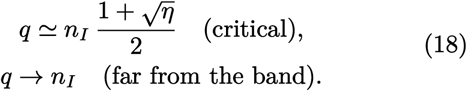

The reporter tail is built during excursions in which LacI is low and the queue is congested, so *q* falls from its critical value to *n*_*I*_ along an excursion. The exponent is then set by an effective *q*_eff_ between the two limits, and Eq. (17) is a finite-window mechanism rather than a universal law. The exponent therefore reports a rate of the sensor gene and the gain of the loop, not the reporter. Simulations confirm Eq. (17) with one fitted *q*_eff_ in place of *q*: the exponent grows with *κ* as predicted until the tail becomes too steep to have a resolvable scaling range [SM Sec. S3, Fig. S2(c)].

Power-law tails are present wherever the loop is in control, and the model delimits this region. In the IPTG-aTc plane, the scaling range is present at every dose aTc while Γ lies in the critical band, and it disappears once IPTG pushes Γ out of it: LacI cannot anymore close the promoter and the reporter distribution collapses to a narrow peak [SM Sec. S3, Figs. S1 and S2(a,b)]. A control in which the LacI message lacks the shRNA target and is therefore blind to the queue still has excursions, since the sensor noise is the same, but around a silenced reporter; reading the queue is what sets the baseline at the critical point instead of at silence. Power laws are thus confined to the region in which the loop is in control.

### D. Measured tails: two reporters, similar exponents

The reporter was measured in two stable HeLa lines carrying the circuit: one with EGFP and one with d2EGFP (Methods). Neither IPTG nor aTc was added during these acquisitions. The data comprise two large flow-cytometry acquisitions, one per construct, with 2.1 million cells in total. Each construct is represented by one clone (SM Sec. S5). The model places this basal, noinducer condition in its predicted critical regime. Thus the measured tails are the predicted untuned phenotype. Because the model was calibrated against the EGFP tail, that reporter’s agreement is by construction; the d2EGFP line provides the comparison across reporter stability.

The heavy tails reflect strong cell-to-cell differences in reporter level that are visible directly in the cells: in a single field of view most cells carry little detectable GFP, while a few neighbors are far brighter (Fig. 4; SM Sec. S5).

**FIG. 4.**
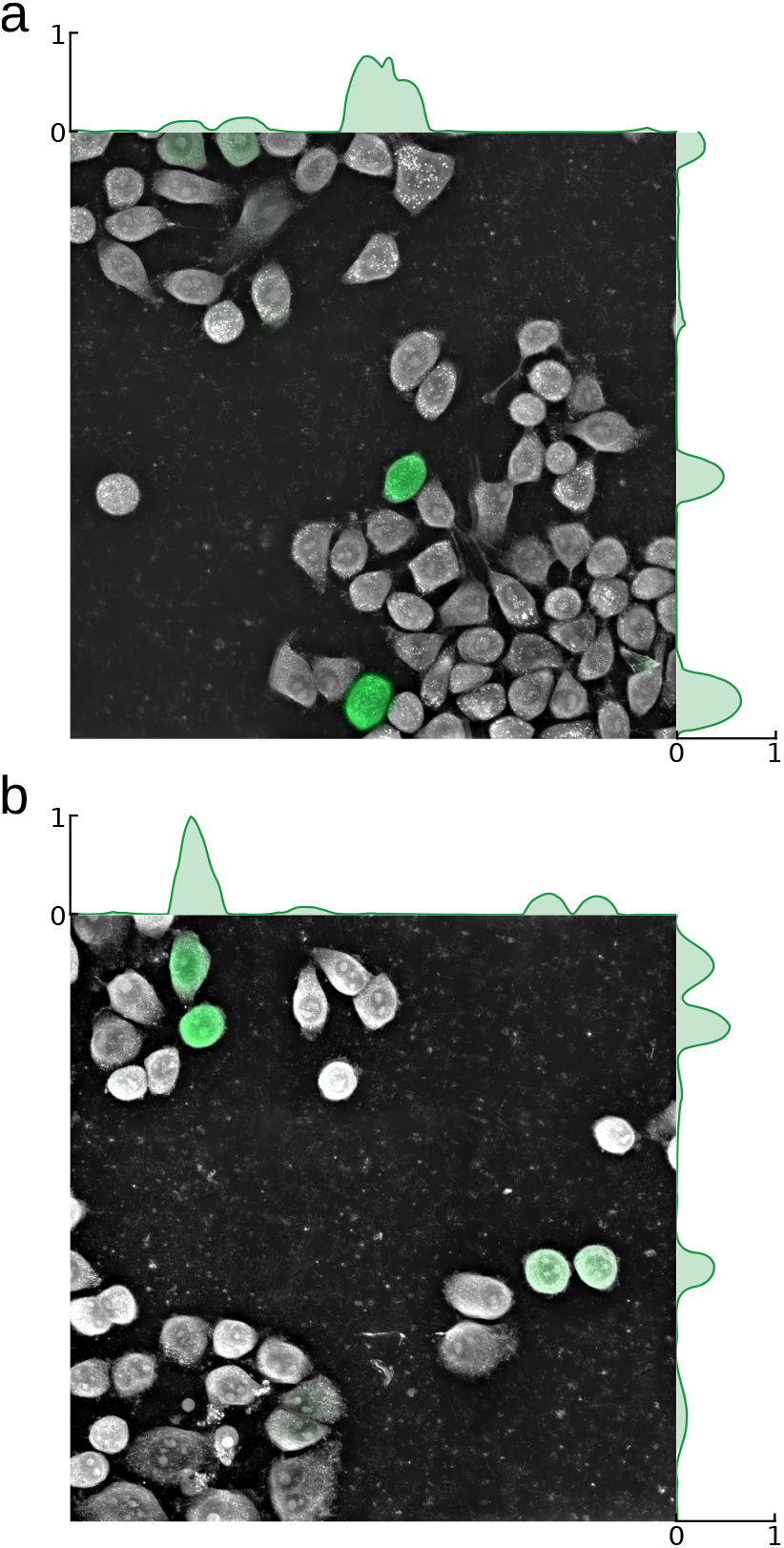
Reporter intensity differs between neighboring cells. Holotomographic images of the (a) d2EGFP and EGFP lines, with no inducer added. Gray, refractive index (maximum-intensity projection); green, background-subtracted GFP fluorescence, on one linear scale for both fields. Curves on the top and right edges are the GFP signal averaged over columns and rows, in units of their common maximum. Most cells are barely fluorescent and a few are bright. Unmerged channels: Fig. S4.

Both distributions exhibit power-law tails spanning more than a decade [Fig. 5(a,b)]. The local exponent, defined from the complementary distribution *P* of the fluorescence divided by its median,

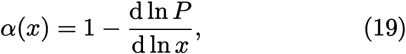

**FIG. 5.**
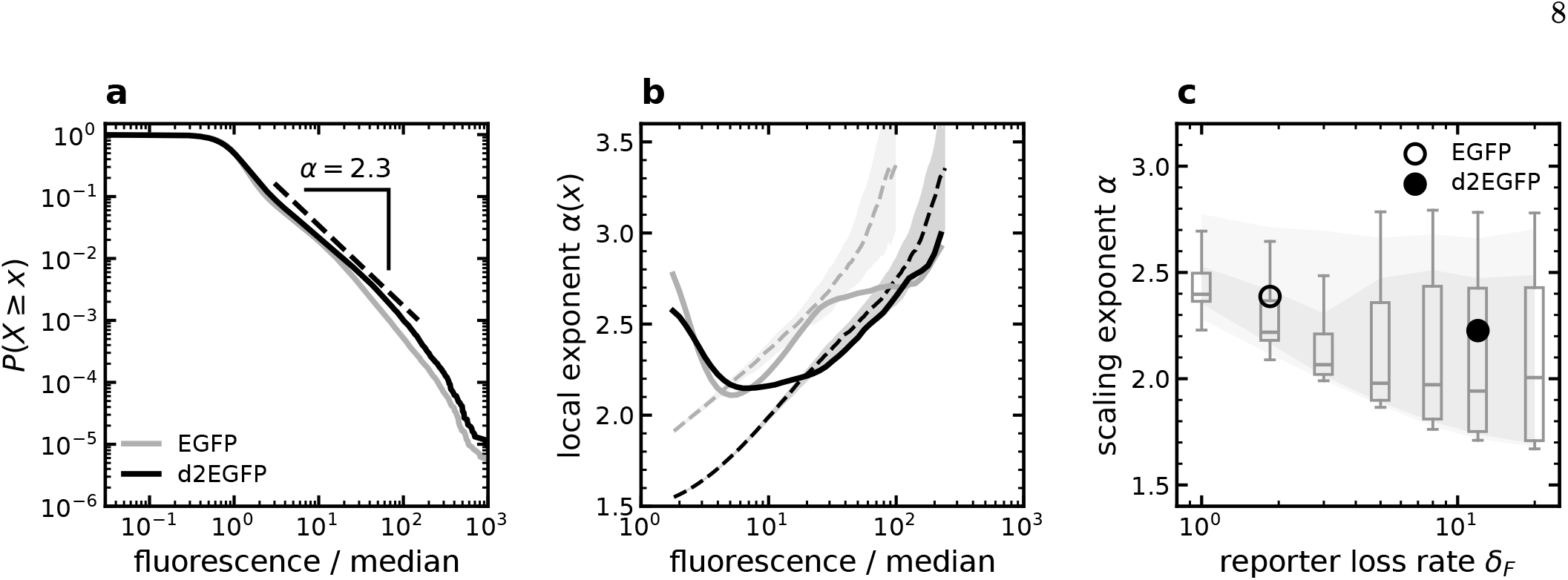
Single-cell fluorescence of the two reporter lines. (a) Complementary fluorescence distributions *P* (*X* ≥ *x*) for EGFP (gray) and d2EGFP (black), each divided by its median (2.1 × 10^6^ cells total). The dashed guide has power-law exponent *α* = 2.3; the inverted L beside it spans one decade in *x* and *α* ™ 1 decades in *P*, the slope of the complementary distribution. (b) Local exponent *α*(*x*) for the same reporter lines; a plateau indicates a power-law range. Thick solid curves are measured; thin dashed curves and shaded bands show model medians and central 68 % intervals across 100 reference-circuit runs, using each reporter’s degradation time. (c) Scaling-range exponent against reporter loss rate *δ*_*F*_, in dilution-rate units. Light gray bands show the central 68 % and 95 % of model runs; gray boxes and whiskers show each run distribution (median, interquartile range, and 1.5-IQR whiskers). The open and filled black circles mark the EGFP and d2EGFP lines, respectively, at their nominal loss rates.

is one plus the steepness of *P* on logarithmic axes. It is constant to within ±0.3 over a range of about 1.5 decades, with the value 2.39 for EGFP and 2.23 for d2EGFP, and steepens beyond this range, as required by a finite cutoff. The criterion is stringent: a lognormal of the same width and sample size has no scaling range of even half a decade.

Conventional maximum-likelihood fits (SM Sec. S5) favor a power law with a finite cutoff over a pure power law, a lognormal or an exponential. A finite cutoff is what the model predicts, because promoter output and queue capacity are finite. The maximum-likelihood exponent depends on the fitting threshold and separates the two lines more than the scaling-range exponent does (SM Sec. S5), so the similarity quoted here refers to the latter.

Four parameters of the model (SM Sec. S1) were calibrated on the tail of the EGFP distribution: the burst frequency *κ* of the sensor gene, the LacI excess Λ, the basal capacity *η*_0_ and the promoter strength *α*_*G*_. The remaining parameters follow from the construct and the literature. The exponent constrains the ratio of *κ* to *q* but not the operating point: Λ, *α*_*G*_ and *η*_0_ enter the tail through Γ and the cutoff and are therefore degenerate, and the acceptable circuits scatter around Γ ≃ 1. That the reference circuit reproduces the EGFP exponent is thus true by construction (Table S2 compares its other tail statistics), and the model is tested instead by the dependence on the reporter.

The reporter comparison supports the model’s prediction that the exponent depends only weakly on reporter lifetime. The tail is generated by excursions of the slow LacI sensor, which last about a dilution lifetime; a reporter that follows the message faster resolves peaks that a slow one smooths away, and therefore has a slightly flatter tail, until it becomes faster than the excursions. The model mean exponent is lower for d2EGFP than for EGFP by 0.19 ± 0.03 (2.28 against 2.09 over 100 runs), and the measured d2EGFP exponent is lower by 0.16. The EGFP exponent was used in calibration, so its agreement is not an independent test; the d2EGFP result is an independent comparison and falls within the model’s run-to-run spread, with the predicted direction and a similar difference. A change of this size corresponds, by Eq. (17), to a change of *κ* of *q*_eff_ Δ*α*, a small fraction of *κ* itself: the exponent is set mainly by the sensor loop and depends only weakly on reporter lifetime.

The model predicts a broader d2EGFP distribution than EGFP with the nominal lifetimes (log-width 1.51 against 1.08; 99th percentile over median 62 against 27), because a short-lived reporter averages the message over a shorter window. The measured d2EGFP distribution is also broader by each scale-free statistic, but the difference is smaller (log-width 0.85 against 0.80; 99th percentile over median 20 against 16; SM Sec. S5). Thus the model captures the direction of the lifetime effect but overestimates its size. The lines are single clones, and the model omits extrinsic variation in copy number and integration site, which can alter the body of each histogram. A second clone per reporter and a decay measurement (SM Sec. S5) would help distinguish clone effects from an incorrect effective d2EGFP loss; neither is expected to change the exponent, which the model fixes mainly through the sensor loop.

## III. DISCUSSION

We have engineered a circuit in mammalian cells in which the shared clearance of messenger RNA acts as a queue with a critical point, and a repressor reads the length of that queue and feeds it back on the demand. In the language of the feedback formulations of SOC [13, 14, 19], the traffic intensity is the control parameter, the queue length is the order parameter, and LacI closes the loop through the lifetime of its own message. In the model, the loop returns the circuit toward the critical point after changes in promoter strength, capacity and inducer dose, provided that the parameter combination Γ remains within a factor of order one of unity; no individual parameter needs to be tuned. The dynamics are those of SOC: slow drive, a threshold and fast relaxation produce intermittent excursions of all sizes with a heavy-tailed distribution and, as in a sandpile, a cutoff set by the finite size of the system, here the capacity of the machinery and the output of the promoter. The scale-free range is therefore of the order of a decade, in the model and in the cells, as expected of critical dynamics in a cell with a finite number of molecules. The critical state is not a fixed point but a state the loop keeps returning to, poised between a free and a congested phase and never settling in either: the “unstable attractor” that the bacterial ClpXP circuit was the first to engineer [61], here in a queue of messenger RNAs instead of proteins.

Three features distinguish this mechanism from other sources of heavy tails, such as mixtures of cell states and multiplicative noise [43, 44]. First, the exponent is a function of the circuit, Eq. (17), so it can be moved by changing the burst frequency of the sensor gene, and it depends on the reporter only weakly, as the two measured lines show. Second, tails exist only where the loop is in control, and the model predicts where they end. Third, the loop makes the reporter follow the capacity that silences it, which repression alone cannot do (SM Sec. S4). Multiplicative noise and mixtures predict neither of the last two. Circuit variants that disable one element at a time are the direct experimental counterparts of the simulated controls: a scrambled target on the *lacI* message makes the sensor blind to the queue; a reporter promoter without *lacO* opens the loop; and a plasmid without the shRNA removes the capacity and with it the queue. The circuit is the mammalian counterpart of the bacterial design. Both use a saturable shared degradation machinery as the queue and a negative feedback that lowers demand when the machinery is congested. The host changes what carries the queue and the noise. Here the queue is RISC-mediated RNA clearance, the sensor is a message whose lifetime lengthens with the queue, and the noise that drives the avalanches is the transcription of a single gene, so that the tail exponent reports the burst frequency *κ* of that gene, which follows the burst frequency and not the message rate if the gene fires in bursts (SM Sec. S3). Bursting kinetics of mammalian genes vary over orders of magnitude [84–86], and *κ* can be measured directly by single-molecule RNA imaging. The bacterial study identified the absence of a temporal dimension as a limitation and proposed that a similar motif in a eukaryotic cell would provide time series. Excursions of the mammalian circuit last about a dilution lifetime, of the order of a cell cycle, so that time-lapse imaging of single-cell lineages over several generations should resolve the intermittency of Fig. 3 directly.

The possible advantages of critical states motivate the search for such circuits in nature. A system poised at a transition has maximal sensitivity and dynamic range, may compute best there [20, 21, 23], and produces broad variability without tuning; broad phenotypic variability is a resource for populations in fluctuating environments [49–51]. If these advantages are real, selection could favor circuits that reach the critical state by feedback rather than by fine tuning [39]. The RNAi machinery is a natural candidate. Endogenous microRNAs and their many targets compete for the same Argonaute pool, and the load of that pool changes between cell types and states [68, 69, 72, 73, 75]. If some endogenous regulators read the load through their own lifetime, as LacI does here, the natural pathway could show the same self-organization; the queue length, its susceptibility and the combination Γ provide a framework for testing this possibility. Analogous queues exist for proteases, proteasomes and ribosomes.

These results also suggest ways to design circuits. A loop that senses a shared, saturable resource can hold expression at a finite-gain operating point despite changes in reporter-specific production and in the capacity of the shared resource [54, 57, 58]. It can also produce controlled, scale-free bursts of expression, with the tail exponent set by one gene’s burst frequency. Such bursts could drive drug delivery or intercellular signaling. Coupling these excursions across cells could produce population-level avalanches.

The measurements analyzed here are from basal conditions, with one clone per reporter. The loss rates for the reporter use nominal values. The model predicts the observed ordering of the distribution widths, but the measured difference is smaller than predicted. Autofluorescence, background subtraction, and extrinsic variation in copy number and cell size [87] are not included in the model and affect the body of histograms more than the tail. A dose matrix across IPTG and aTc, perturbations of capacity and load (SM Sec. S4), and the feedback-disrupted variants described above will test the model’s predictions using the same lines and detector.

Beyond its role as a minimal realization of SOC, the mammalian circuit opens several possibilities for engineering collective dynamics in synthetic multicellular systems. One particularly interesting direction concerns the emergence of collective sensing, memory and computation in cellular assemblies [88]. By coupling the intermittent expression bursts generated by individual SOC circuits through intercellular signaling, one could explore whether local fluctuations become coordinated into collective avalanches, and whether proximity to criticality enhances the sensitivity, dynamic range and information-processing capabilities of the resulting tissue. Embedding such circuits in synthetic aggregates, spheroids or organoids would provide a controlled platform to investigate how cell-autonomous regulatory dynamics contribute to higher-level forms of organization and basal cognition. More generally, the possibility of engineering critical dynamics within living cells offers a new route to interrogate the physical principles underlying multicellular complexity: rather than merely searching for signatures of criticality in natural tissues, we can construct the mechanisms that generate them and test whether criticality provides a functional advantage for collective behavior, adaptation and information processing.

## IV. MATERIALS AND METHODS

### A. Circuit and cell lines

A single plasmid carries four transcription units: CMV-*lacI-NLS* followed by an shRNA target site; U6-tetO expressing the target-specific shRNA; CMV-*tetR*; and RSV-*lacO*_3_ driving EGFP or d2EGFP [89] followed by the same target site. aTc inhibits TetR and thus derepresses the shRNA; IPTG inhibits LacI. The two reporter versions differ only in the d2 degron. Constructs were assembled by scarless *in vivo* assembly in *E. coli* [90] from PCR fragments with 15-20-nt homology overhangs; products were DpnI-treated, colonies were screened across the junctions, and junctions and whole plasmids were verified by Sanger and whole-plasmid sequencing.

HeLa cells (ATCC CCL-2) were maintained in DMEM with 10 % FBS and 1 % penicillin-streptomycin at 37 ^°^C and 5 % CO_2_, passaged below 80 % confluence and screened monthly for mycoplasma. For stable lines, 2.5 × 10^5^ cells per well were seeded in six-well plates 24 h before transfection with Lipofectamine 3000. Hygromycin selection began 24 h after transfection at 200 µg mL^™1^ and continued for 12 d; the maintenance concentration thereafter was 100 µg mL^™1^. Cells were first sorted for high GFP with 10 mmol L^™1^ IPTG. After 72 h without inducer, they were sorted again for low GFP, and single cells were deposited into 96-well plates with conditioned medium using debris, singlet and viability gates. This selected cells with IPTG-dependent reporter expression. Wells with more than one focus were discarded and clones were expanded from single foci.

### B. Flow cytometry

Cells were detached, resuspended in PBS with 2 % FBS and analyzed on a BD LSRFortessa flow cytometer. GFP was measured with 488 nm excitation using a 505 nm long-pass filter and a band-pass filter centered at 530 nm with a 28 nm bandwidth. The two acquisitions analyzed here are one sample per HeLa construct; neither IPTG nor aTc was added. The exports contain background-subtracted intensities, but do not include acquisition gates, session identifiers, full instrument settings or the background-subtraction procedure.

### C. Tail analysis

The same procedure is applied to measured and simulated distributions. The local exponent of Eq. (19) is computed by least squares over ±0.25 decades wherever at least 40 events lie above *x*. The scaling range is the longest interval on which *α*(*x*) stays within ±0.3 of its median, of at least half a decade, and the exponent of the range is that median. As a conventional cross-check, a Pareto tail is fitted by maximum likelihood above a Kolmogorov-Smirnov-selected threshold [91, 92] and compared with a tapered power law, a truncated lognormal and an exponential (SM Sec. S5).

### D. Model and simulation

The messages and LacI protein are simulated as a stochastic reaction network coupled through shared message clearance (Fig. 1; SM Sec. S1). Time is measured in dilution lifetimes and message concentrations in units of the RISC Michaelis constant. Gillespie’s algorithm simulates these reactions exactly [83]. Reporter protein is represented by the conditional mean of its production and loss process, which is advanced exactly between reaction events; intrinsic reporter-protein birth and death noise is not sampled. Deterministic fixed points are those of the mean-field limit of the same model. Controls remove one element: reporter-promoter repression (open loop) or the target site of the *lacI* message (blind sensor). Every assumption has a biological reading, given in SM Sec. S1, and only four parameters were calibrated on the data.

## ACKNOWLEDGMENTS

The authors thank members of the Complex Systems Lab for valuable discussions and the Santa Fe Institute. Special thanks to Victor Maull for early work on this project. We also thank the UPF Flow Cytometry Core Facility for its support and access to the BD LSRFortessa. Holotomographic images were acquired at the Mesoscopic Imaging Facility of EMBL, which we thank for its support. This work was supported by AGAUR FI-SDUR 2020, grant AEI-PID2023-152129NB-I00, and the Santa Fe Institute. JPM was supported by grant PRE2020-091968, funded by MCIN/AEI (10.13039/501100011033) and co-funded by the European Social Fund through the program *Investing n your future*. NCP was supported by an MCIN/AEI postdoctoral award (reference CEX2018-000792-M).

## References

[1] H. E. Stanley, Reviews of Modern Physics 71, S358 (1999).

[2] J. P. Sethna, K. A. Dahmen, and C. R. Myers, Nature 410, 242 (2001).

[3] N. Goldenfeld, Lectures on phase transitions and the renormalization group (CRC Press, 2018).

[4] P. Bak, C. Tang, and K. Wiesenfeld, Physical Review Letters 59, 381 (1987).

[5] P. Bak, C. Tang, and K. Wiesenfeld, Physical Review A 38, 364 (1988).

[6] V. Frette, K. Christensen, A. Malthe-Sørenssen, J. Feder, T. Jøssang, and P. Meakin, Nature 379, 49 (1996).

[7] L. A. N. Amaral and K. B. Lauritsen, Physical Review E 54, R4512 (1996).

[8] B. Drossel and F. Schwabl, Physical Review Letters 69, 1629 (1992).

[9] R. V. Solé, D. Alonso, and A. McKane, Philosophical Transactions of the Royal Society B: Biological Sciences 357, 667 (2002).

[10] R. V. Solé and S. C. Manrubia, Journal of Theoretical Biology 173, 31 (1995).

[11] P. Bak and K. Sneppen, Physical Review Letters 71, 4083 (1993).

[12] R. V. Solé, S. C. Manrubia, M. Benton, S. Kauffman, and P. Bak, Trends in ecology & evolution 14, 156 (1999).

[13] D. Sornette, Journal de Physique I 2, 2065 (1992).

[14] R. Dickman, A. Vespignani, and S. Zapperi, Physical Review E 57, 5095 (1998).

[15] D. Dhar, Physica A: Statistical Mechanics and its Applications 263, 4 (1999).

[16] R. Dickman, M. A. Munoz, A. Vespignani, and S. Zapperi, Brazilian Journal of Physics 30, 27 (2000).

[17] J. A. Bonachela and M. A. Muñoz, Journal of Statistical Mechanics: Theory and Experiment 2009, P09009 (2009).

[18] G. Pruessner, Self-Organised Criticality: Theory, Models and Characterisation (Cambridge University Press, Cambridge, 2012).

[19] V. Buendía, S. di Santo, J. A. Bonachela, and M. A. Muñoz, Frontiers in Physics 8, 333 (2020).

[20] O. Kinouchi and M. Copelli, Nature Physics 2, 348 (2006).

[21] W. L. Shew, H. Yang, T. Petermann, R. Roy, and D. Plenz, Journal of Neuroscience 29, 15595 (2009).

[22] J. M. Beggs, Philosophical Transactions of the Royal Society A: Mathematical, Physical and Engineering Sciences 366, 329 (2008).

[23] C. G. Langton, Physica D: Nonlinear Phenomena 42, 12 (1990).

[24] J. M. Beggs and D. Plenz, Journal of Neuroscience 23, 11167 (2003).

[25] D. R. Chialvo, Nature Physics 6, 744 (2010).

[26] W. L. Shew and D. Plenz, The Neuroscientist 19, 88 (2013).

[27] J. Hesse and T. Gross, Frontiers in Systems Neuroscience 8, 166 (2014).

[28] A. Cavagna, A. Cimarelli, I. Giardina, G. Parisi, R. Santagati, F. Stefanini, and M. Viale, Proceedings of the National Academy of Sciences 107, 11865 (2010).

[29] W. Bialek, A. Cavagna, I. Giardina, T. Mora, E. Silvestri, M. Viale, and A. M. Walczak, Proceedings of the National Academy of Sciences 109, 4786 (2012).

[30] M. Takayasu, K. Fukuda, and H. Takayasu, Physica A: Statistical Mechanics and its Applications 274, 140 (1999).

[31] R. V. Solé and S. Valverde, Physica A: Statistical Mechanics and its Applications 289, 595 (2001).

[32] K. Nagel and M. Paczuski, Physical Review E 51, 2909 (1995).

[33] T. Nagatani, Reports on progress in physics 65, 1331 (2002).

[34] S. A. Kauffman, Journal of Theoretical Biology 22, 437 (1969).

[35] E. Balleza, E. R. Alvarez-Buylla, A. Chaos, S. Kauffman, I. Shmulevich, and M. Aldana, PLoS ONE 3, e2456 (2008).

[36] M. Nykter, N. D. Price, M. Aldana, S. A. Ramsey, S. A. Kauffman, L. E. Hood, O. Yli-Harja, and I. Shmulevich, Proceedings of the National Academy of Sciences 105, 1897 (2008).

[37] S. Bornholdt and T. Rohlf, Physical Review Letters 84, 6114 (2000).

[38] C. Torres-Sosa, S. Huang, and M. Aldana, PLoS Computational Biology 8, e1002669 (2012).

[39] J. Hidalgo, J. Grilli, S. Suweis, M. A. Muñoz, J. R. Banavar, and A. Maritan, Proceedings of the National Academy of Sciences 111, 10095 (2014).

[40] T. Mora and W. Bialek, Journal of Statistical Physics 144, 268 (2011).

[41] M. A. Munóz, Reviews of Modern Physics 90, 031001 (2018).

[42] M. E. J. Newman, Contemporary Physics 46, 323 (2005).

[43] M. P. H. Stumpf and M. A. Porter, Science 335, 665 (2012).

[44] J. Touboul and A. Destexhe, Physical Review E 95, 012413 (2017).

[45] V. Priesemann, M. Wibral, M. Valderrama, R. Pröpper, M. Le Van Quyen, T. Geisel, J. Triesch, D. Nikolić, and M. H. J. Munk, Frontiers in Systems Neuroscience 8, 108 (2014).

[46] M. B. Elowitz, A. J. Levine, E. D. Siggia, and P. S. Swain, Science 297, 1183 (2002).

[47] A. Raj and A. van Oudenaarden, Cell 135, 216 (2008).

[48] A. Eldar and M. B. Elowitz, Nature 467, 167 (2010).

[49] E. Kussell and S. Leibler, Science 309, 2075 (2005).

[50] N. Q. Balaban, J. Merrin, R. Chait, L. Kowalik, and S. Leibler, Science 305, 1622 (2004).

[51] J.-W. Veening, W. K. Smits, and O. P. Kuipers, Annual Review of Microbiology 62, 193 (2008).

[52] M. B. Elowitz and S. Leibler, Nature 403, 335 (2000).

[53] S. Hooshangi, S. Thiberge, and R. Weiss, Proceedings of the National Academy of Sciences 102, 3581 (2005).

[54] A. Gyorgy, J. I. Jiménez, J. Yazbek, H.-H. Huang, H. Chung, R. Weiss, and D. Del Vecchio, Biophysical Journal 109, 639 (2015).

[55] Y. Qian, H.-H. Huang, J. I. Jiménez, and D. Del Vecchio, ACS Synthetic Biology 6, 1263 (2017).

[56] D. Del Vecchio, A. J. Dy, and Y. Qian, Journal of the Royal Society Interface 13, 20160380 (2016).

[57] T. Frei, F. Cella, F. Tedeschi, J. Gutiérrez, G.-B. Stan, M. Khammash, and V. Siciliano, Nature Communications 11, 4641 (2020).

[58] F. Ceroni, A. Boo, S. Furini, T. E. Gorochowski, O. Borkowski, Y. N. Ladak, A. R. Awan, C. Gilbert, G.-B. Stan, and T. Ellis, Nature Methods 15, 387 (2018).

[59] R. D. Jones, Y. Qian, V. Siciliano, B. DiAndreth, J. Huh, R. Weiss, and D. Del Vecchio, Nature Communications 11, 5690 (2020).

[60] N. A. Cookson, W. H. Mather, T. Danino, O. Mondragón-Palomino, R. J. Williams, L. S. Tsimring, and J. Hasty, Molecular Systems Biology 7, 561 (2011).

[61] B. Vidiella, A. Guillamon, J. Sardanyés, V. Maull, J. Pla, N. Conde, and R. Solé, Nature Communications 12, 4415 (2021).

[62] A. Fire, S. Xu, M. K. Montgomery, S. A. Kostas, S. E. Driver, and C. C. Mello, Nature 391, 806 (1998).

[63] S. M. Elbashir, J. Harborth, W. Lendeckel, A. Yalcin, K. Weber, and T. Tuschl, Nature 411, 494 (2001).

[64] G. Hutvágner and P. D. Zamore, Science 297, 2056 (2002).

[65] D. P. Bartel, Cell 136, 215 (2009).

[66] M. Ghildiyal and P. D. Zamore, Nature Reviews Genetics 10, 94 (2009).

[67] S. Jonas and E. Izaurralde, Nature Reviews Genetics 16, 421 (2015).

[68] A. Arvey, E. Larsson, C. Sander, C. S. Leslie, and D. S. Marks, Molecular Systems Biology 6, 363 (2010).

[69] A. A. Khan, D. Betel, M. L. Miller, C. Sander, C. S. Leslie, and D. S. Marks, Nature Biotechnology 27, 549 (2009).

[70] S. Mukherji, M. S. Ebert, G. X. Y. Zheng, J. S. Tsang, P. A. Sharp, and A. van Oudenaarden, Nature Genetics 43, 854 (2011).

[71] D. Grimm, K. L. Streetz, C. L. Jopling, T. A. Storm, K. Pandey, C. R. Davis, P. Marion, F. Salazar, and M. A. Kay, Nature 441, 537 (2006).

[72] L. Salmena, L. Poliseno, Y. Tay, L. Kats, and P. P. Pandolfi, Cell 146, 353 (2011).

[73] C. Bosia, A. Pagnani, and R. Zecchina, PLoS ONE 8, e66609 (2013).

[74] M. Figliuzzi, E. Marinari, and A. De Martino, Biophysical Journal 104, 1203 (2013).

[75] R. Denzler, V. Agarwal, J. Stefano, D. P. Bartel, and M. Stoffel, Molecular Cell 54, 766 (2014).

[76] U. Ala, F. A. Karreth, C. Bosia, A. Pagnani, R. Taulli, V. Léopold, Y. Tay, P. Provero, R. Zecchina, and P. P. Pandolfi, Proceedings of the National Academy of Sciences 110, 7154 (2013).

[77] M. Gossen and H. Bujard, Proceedings of the National Academy of Sciences USA 89, 5547 (1992).

[78] M. van de Wetering, I. Oving, V. Muncan, M. T. Pon Fong, H. Brantjes, D. van Leenen, F. C. P. Holstege, T. R. Brummelkamp, R. Agami, and H. Clevers, EMBO Reports 4, 609 (2003).

[79] T. L. Deans, C. R. Cantor, and J. J. Collins, Cell 130, 363 (2007).

[80] J. D. C. Little, Operations Research 9, 383 (1961).

[81] S. Oehler, E. R. Eismann, H. Krämer, and B. Müller-Hill, The EMBO Journal 9, 973 (1990).

[82] T. Kuhlman, Z. Zhang, M. H. Saier, and T. Hwa, Proceedings of the National Academy of Sciences 104, 6043 (2007).

[83] D. T. Gillespie, The Journal of Physical Chemistry 81, 2340 (1977).

[84] A. Raj, C. S. Peskin, D. Tranchina, D. Y. Vargas, and S. Tyagi, PLoS Biology 4, e309 (2006).

[85] D. M. Suter, N. Molina, D. Gatfield, K. Schneider, U. Schibler, and F. Naef, Science 332, 472 (2011).

[86] A. J. M. Larsson, P. Johnsson, M. Hagemann-Jensen, L. Hartmanis, O. R. Faridani, B. Reinius, Å. Segerstolpe, C. M. Rivera, B. Ren, and R. Sandberg, Nature 565, 251 (2019).

[87] O. Padovan-Merhar, G. P. Nair, A. G. Biaesch, A. Mayer, S. Scarfone, S. W. Foley, A. R. Wu, L. S. Churchman, A. Singh, and A. Raj, Molecular Cell 58, 339 (2015).

[88] R. Solé, N. Conde-Pueyo, J. Pla-Mauri, J. Garcia-Ojalvo, N. Montserrat, and M. Levin, NPJ systems biology and applications 10, 151 (2024).

[89] X. Li, X. Zhao, Y. Fang, X. Jiang, T. Duong, C. Fan, C.-C. Huang, and S. R. Kain, Journal of Biological Chemistry 273, 34970 (1998).

[90] J. García-Nafría, J. F. Watson, and I. H. Greger, Scientific Reports 6, 27459 (2016).

[91] A. Clauset, C. R. Shalizi, and M. E. J. Newman, SIAM Review 51, 661 (2009).

[92] J. Alstott, E. Bullmore, and D. Plenz, PLoS ONE 9, e85777 (2014).

