## Supplementary Material for "Self-organized criticality in a mammalian RNA-interference circuit"

### Supplementary Material
### Self-organized criticality in a mammalian RNA-interference circuit


Jordi Pla-Mauri,[1,2,*] Marc Güell,[3,4] Ricard Solé,[1,2,4,5,†] and Núria Conde-Pueyo[1,6,‡]

[1]*Complex Systems Lab, Universitat Pompeu Fabra, Dr. Aiguader 88, 08003 Barcelona, Spain*
[2]*Institut de Biologia Evolutiva, CSIC-UPF, Pg. Marítim de la Barceloneta 37, 08003 Barcelona, Spain*
[3]*Department of Medicine and Life Sciences, Universitat Pompeu Fabra, 08003 Barcelona, Spain*
[4]*Institució Catalana de Recerca i Estudis Avançats (ICREA), Pg. Lluís Companys 23, 08010 Barcelona, Spain*
[5]*Santa Fe Institute, 1399 Hyde Park Road, Santa Fe, New Mexico 87501, USA*
[6]*European Molecular Biology Laboratory, EMBL Barcelona, Dr. Aiguader 88, 08003 Barcelona, Spain*


#### S1. MODEL AND SIMULATION

##### A. Reaction network

The stochastic state variables are the numbers of *lacI* messages $n_{I_m}$, reporter messages $n_{G_m}$ and LacI protein $n_{\mathrm{p}}$. Reporter protein is treated separately as an output filter; TetR and the shRNA enter through the capacity $\eta(r)$. With $\Omega$ the Michaelis constant expressed as a number of target molecules, the definitions are

$$M = \frac{n_{I_m} + n_{G_m}}{\Omega}, \qquad I = \frac{n_{\mathrm{p}}}{\Omega},$$

$$\varepsilon = \frac{\delta_c}{\delta_m}, \qquad r = \frac{[\mathrm{aTc}]}{K_r},$$

$$\eta(r) = \eta_0 + (\eta_\infty - \eta_0)\,\frac{r}{1+r},$$

$$\Theta = \frac{\Lambda}{\alpha_I}, \qquad \kappa = \frac{\Omega\alpha_I}{\varepsilon\, b_I},$$

$$\eta - 1 = \frac{k_{\mathrm{cat}} R_{\mathrm{tot}}}{K_m \delta_m}.$$

Here $M$ is the total load, $I$ is in units of $K_L$ (counted on the same scale $\Omega$ as $M$), $\varepsilon$ compares the dilution rate with the basal message loss rate, $\Theta$ is the translation rate ($\beta_I$ of the main text, in units of $K_L$), $\kappa$ is the frequency at which the *lacI* gene fires per dilution lifetime, and the capacity is the catalytic rate times the RISC pool over the Michaelis constant and the basal message loss rate. The reactions are

$$\varnothing \to I_m \qquad \text{rate } \frac{\Omega\alpha_I}{\varepsilon b_I},$$

$$\varnothing \to G_m \qquad \text{rate } \frac{\Omega\alpha_G H(I)}{\varepsilon b_G},$$

$$I_m \to \varnothing \qquad \text{rate } \frac{n_{I_m}\,\phi(M)}{\varepsilon},$$

$$G_m \to \varnothing \qquad \text{rate } \frac{n_{G_m}\,\phi(M)}{\varepsilon},$$

$$I_m \to I_m + I_{\mathrm{p}} \qquad \text{rate } \Theta n_{I_m},$$

$$I_{\mathrm{p}} \to \varnothing \qquad \text{rate } n_{\mathrm{p}},$$

where each birth adds a geometric burst of mean size $b_I$ or $b_G$ messages and $H$ is the promoter activity of Eq. (9). Reporter protein follows

$$\frac{\mathrm{d}F}{\mathrm{d}t} = k_F\, G_m - \delta_F\, F \tag{S1}$$

| Parameter | Meaning | Value | Basis |
|---|---|---|---|
| $\varepsilon = \delta_c/\delta_m$ | dilution over message loss | 0.1 | literature |
| $\Omega$ | $K_m$ in molecules (counting scale) | 10 | insensitive |
| $\kappa = \Omega\alpha_I/(\varepsilon b_I)$ | LacI gene bursts per dilution lifetime | 3.3 | tail |
| $b_I$, $b_G$ | mean burst size of the LacI and reporter genes | 1, 1 | reference (varied in Table S3) |
| $\Lambda = \beta_I\alpha_I/K_L$ | LacI excess over operator affinity | 57 | tail (degenerate) |
| $n_I$ | repression cooperativity | 2 | construct (LacI loops between two operators) |
| $\eta_0$ | knock-down fold, no aTc | 24 | tail |
| $\eta_\infty$ | knock-down fold, saturating aTc | 100 | nominal |
| $K_r$, $n_r$, $K_i$ | aTc and IPTG dose scales, aTc Hill exponent | 1, 1, 1 | nominal |
| $\alpha_G$ | reporter promoter output | 2600 | tail |
| $\Gamma_0$ | $\Lambda/(\alpha_G^{1/n_I}\eta_0^{1/2-1/n_I})$: operating point at zero inducers | 1.12 | derived |
| $\delta_F$ | reporter loss, EGFP / d2EGFP | 1.85 / 12 | literature |

TABLE S1. **Reference parameters.** Time is in dilution lifetimes and concentrations in units of $K_m$. "Construct" and "literature" values are fixed a priori; "tail" values were adjusted to the tail quantiles of the EGFP distribution; dose-response parameters are nominal values for the model; "derived" values follow from the others.

with loss rate $\delta_F$. Conditional on the message path, reporter protein follows an immigration-death process. The simulations advance its conditional mean exactly between reaction events and do not sample intrinsic reporter-protein birth and death noise; $k_F$ sets only the units.

##### B. Biological reading of the assumptions

The model represents the two messages as substrates of one enzyme with one Michaelis constant because they carry the same target sequence. This is a simplifying assumption about equal effective clearance, not a measured property of the two RNAs; target context, localization and pathway kinetics could make their clearance differ. The silencing capacity is a mean over the shRNA pool, which is large and long-lived compared with a message, and aTc enters only through that mean. LacI is a stable nuclear protein, so its loss is dilution alone. The repression cooperativity $n_I = 2$ reflects the two DNA-binding heads of the LacI tetramer, which loop between two operators of the reporter promoter. Each transcription event of the *lacI* gene makes one message in the reference ($b_I = 1$) and a burst in Table S3; the reporter gene is Poisson in the reference and bursty in Table S3. Plasmid copy number, cell size and the cell cycle are constant in the model, so extrinsic noise, which the measured histograms contain, is not represented.

##### C. Parameters and calibration

Table S1 lists the parameters. Literature values are $\varepsilon$ (a message lifetime of a few hours at a dilution lifetime of about 1 d), the reporter loss rates

$$\delta_F = 1 + \frac{t_d}{t_{1/2}} \tag{S2}$$

from the degradation half-lives $t_{1/2}$ of EGFP [1] and d2EGFP [2] at the doubling time $t_d$, and $n_I$ from the construct. The burst frequency $\kappa$, the LacI excess $\Lambda$, the basal capacity $\eta_0$ and the promoter strength $\alpha_G$ were adjusted in two stages. Stage one is a random search over 192 circuits, log-uniform in each parameter, with one run each; stage two reruns the 8 best circuits several times for longer. Both stages minimize the score

$$\sum_k \ln^2 \frac{q_k^{\text{model}}}{q_k^{\text{data}}} + 4\,(\alpha - \alpha_{\text{data}})^2, \tag{S3}$$

where $q_k$ are the 99th, 99.9th and 99.99th percentiles of the EGFP distribution (divided by the median) and $\alpha$ is the scaling exponent, against the measured EGFP line; the body width is not scored because autofluorescence and extrinsic variation, which the model omits, dominate it. Only 8 of the 192 circuits are acceptable, and they scatter around $\Gamma \simeq 1$. The best finalist, rounded to two significant figures, defines the reference circuit; its exponent matches the measured EGFP exponent, and it is rescored on independent long runs. The exponent of the scaling range of a single run scatters through the choice of window (standard deviation 0.15 for the reference circuit), so every model

| | EGFP | d2EGFP | model |
|---|---|---|---|
| positive events | 1 437 041 | 677 988 | |
| median (a.u.) | 69 | 90 | |
| mean / median | 1.77 | 2.02 | 2.61 |
| 99th pct. / median | 16.3 | 19.6 | 27.2 |
| 99.9th pct. / median | 69 | 101 | 91 |
| log-width $\sigma_{\log}$ | 0.80 | 0.85 | 1.08 |
| scaling exponent $\alpha$ | 2.39 | 2.23 | 2.28 |
| range ($\times$ median) | 2.0–56 | 2.2–63 | 3.0–31 |
| range (decades) | 1.45 | 1.45 | 0.96 |
| Pareto exponent | 2.68 | 2.19 | |
| threshold / median | 25.8 | 3.3 | |
| $\Delta$AIC, Pareto | 11 | 209 | |
| $\Delta$AIC, lognormal | 5 | 116 | |
| $\Delta$AIC, exponential | 2778 | 43390 | |
| cutoff $x_c/x_{\min}$ | 55 | 152 | |
| Pareto GOF $p$ | 0.01 | 0.07 | |
| non-positive | 0.40 % | 0.23 % | |

TABLE S2. **The two reporter distributions.** Shape statistics are divided by the median. The scaling exponent is the median local exponent over the scaling range ("range"); the Pareto exponent is the maximum-likelihood fit above the Kolmogorov-Smirnov-selected threshold. $\Delta$AIC is relative to the best model on the same tail support (the tapered power law), whose cutoff is $x_c$; GOF is the goodness-of-fit test on 5000 events. The last column is the reference circuit with the EGFP filter.

exponent quoted is a mean over runs. The dose scales $K_i$, $K_r$ and the maximal capacity $\eta_\infty$ are nominal values for these model calculations; dose-response measurements will set their assay-specific values. The number $\Omega$ is a counting scale, and the shapes reported are insensitive to it (Table S3). The free, critical and congested cases of Fig. 3 are model settings, not separate measurements: the free case reduces $\alpha_G$ a hundredfold, while the congested case sets $i = 30$. At the reference reporter strength the free phase cannot be reached with nonnegative IPTG; the free trajectory therefore represents a weaker-reporter model setting rather than an inducer-dose condition.

##### D. Validation

The simulator reproduces the exact stationary distribution of the frozen-input queue, a confluent-hypergeometric law that follows from detailed balance, and the exact mean of the reporter filter; its deterministic limits reproduce the closed-form identities of the main text.

#### S2. DERIVATIONS

*a. Susceptibility and critical band.* The balance $A = M\phi(M)$ gives

$$A = \frac{M(M+\eta)}{M+1}, \qquad \frac{\mathrm{d}A}{\mathrm{d}M} = \frac{M^2+2M+\eta}{(M+1)^2}, \tag{S4}$$

and, by Eq. (5), the susceptibility is

$$S_A = \frac{\mathrm{d}\ln M}{\mathrm{d}\ln A} - 1 = \frac{(\eta-1)M}{M^2+2M+\eta}. \tag{S5}$$

It is maximal at $M^* = \sqrt{\eta}$, which is exactly $\rho = 1$, where $S_A = S_{\max}$ of Eq. (7). The half-maximum condition is the quadratic $M^2 - (4\sqrt{\eta}+2)M + \eta = 0$, with roots

$$M_\pm = 2\sqrt{\eta} + 1 \pm \sqrt{3\eta + 4\sqrt{\eta} + 1} \simeq (2 \pm \sqrt{3})\sqrt{\eta}. \tag{S6}$$

A load $M = c\sqrt{\eta}$ corresponds to

$$\rho \simeq 1 + \frac{c - 1/c}{\sqrt{\eta}}, \tag{S7}$$

so the band edges are those of Eq. (7).

*b. Set point.* For a clearance rate $\phi$ the load and the demand are

$$M = \frac{\eta - \phi}{\phi - 1}, \qquad A = M\phi, \tag{S8}$$

which gives Eq. (12) exactly. For $\phi = x\sqrt{\eta}$ and large $\eta$,

$$\rho \simeq \Big(1 + \frac{1}{x\sqrt{\eta}}\Big)\Big(1 - \frac{x}{\sqrt{\eta}}\Big) \simeq 1 + \frac{x^{-1} - x}{\sqrt{\eta}}. \tag{S9}$$

The edges of the critical band of Eq. (7) correspond to $x^{-1} - x = \pm 2\sqrt{3}$, that is, to $x = 2 - \sqrt{3}$ (upper edge) and $x = 2 + \sqrt{3}$ (lower edge). With the reporter carrying nearly all the demand, $A_G \simeq \rho\eta$, and

$$H \simeq \frac{\rho\eta}{\alpha_G}, \qquad D \simeq \frac{\alpha_G}{\rho\eta}, \tag{S10}$$

so at $\rho \simeq 1$ Eq. (11) gives

$$\frac{\phi^*}{\sqrt{\eta}} \simeq \Gamma. \tag{S11}$$

*c. Loop gain and compression.* Let the reporter promoter strength be $\lambda\alpha_G$. Then $\mathrm{d}\ln I = S_A\,\mathrm{d}\ln A$ from Eq. (8), $\mathrm{d}\ln H = -n_I(1-H)\,\mathrm{d}\ln I$ from the promoter, and $\mathrm{d}A = \mathrm{d}A_G$ with $A_G = \lambda\alpha_G H$. Combining these,

$$\frac{\mathrm{d}\ln A^*}{\mathrm{d}\ln\lambda} = \frac{A_G/A}{1+\mathcal{L}}, \qquad \mathcal{L} = hS_A, \quad h = n_I(1-H)\frac{A_G}{A}. \tag{S12}$$

At the critical point $h \simeq n_I$ and $S_A = S_{\mathrm{max}}$, so

$$\mathcal{L} \simeq n_I\,\frac{\sqrt{\eta} - 1}{2}, \tag{S13}$$

and the feedback compresses a change of promoter strength by the loop gain, which is largest inside the critical band because $S_A$ is.

*d. Tail exponent.* The reporter message is $G^* = A_G W$. Using

$$\mathrm{d}\ln A_G = -n_I(1-H)\,\mathrm{d}\ln I, \quad \mathrm{d}\ln A = \frac{A_G}{A}\,\mathrm{d}\ln A_G, \quad \mathrm{d}\ln W = S_A\,\mathrm{d}\ln A, \tag{S14}$$

one finds

$$\mathrm{d}\ln G^* = -n_I(1-H)\Big[1 + \frac{A_G}{A}S_A\Big]\mathrm{d}\ln I, \tag{S15}$$

which is Eq. (16). If $P(I < x) \propto x^{\kappa}$ at low $I$, then

$$P(G > g) = P\big(I < g^{-1/q}\big) \propto g^{-\kappa/q}, \tag{S16}$$

so the density has the exponent of Eq. (17).

#### S3. WHERE THE POWER LAWS LIVE

Figure S1 shows the closed-loop reporter distribution for increasing IPTG. At low doses the distributions share a scale-free body; at high doses the promoter opens, the traffic intensity leaves the critical band and the distribution collapses to a narrow peak. The open loop, matched to the same deterministic demand at the reference condition, lacks this response across the dose range. Figure S2(a,b) maps the scaling range and exponent over the IPTG-aTc plane. At every aTc dose the tail persists for IPTG up to about the dose at which $\Gamma$ leaves the critical band, $1+i \simeq (2+\sqrt{3})\Gamma_0$, and it is absent beyond; the normalized distributions are similar at all doses even though the median reporter level changes with aTc (Sec. S4). Figure S2(c) tests Eq. (17): the exponent grows with $\kappa$ with an effective $q_{\mathrm{eff}}$ between $n_I$, its value far from the critical band, and its value at the critical point [Eq. (18)], since $q$ falls from the second value toward the first along an excursion.

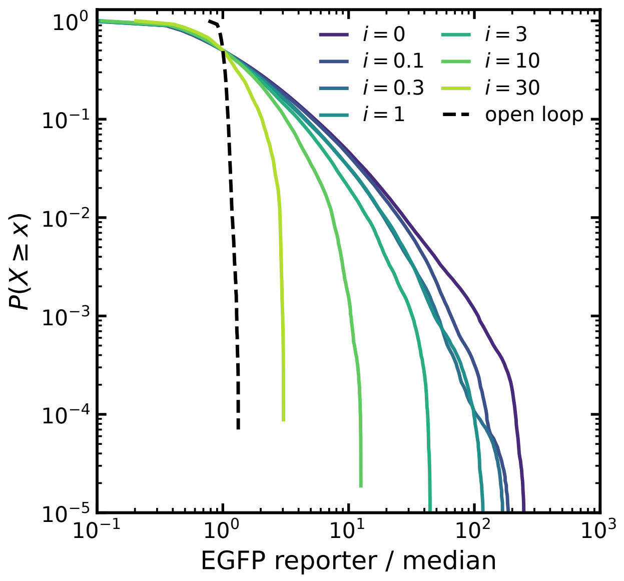


FIG. S1. **Reporter distributions against** IPTG. Complementary distributions of the EGFP reporter (divided by its median) in the closed loop at zero aTc for increasing IPTG dose $i$ (from dark to light), and for the open loop at the same deterministic demand at the reference condition (black dashed). Gillespie simulations with reporter protein represented by its conditional mean filter.

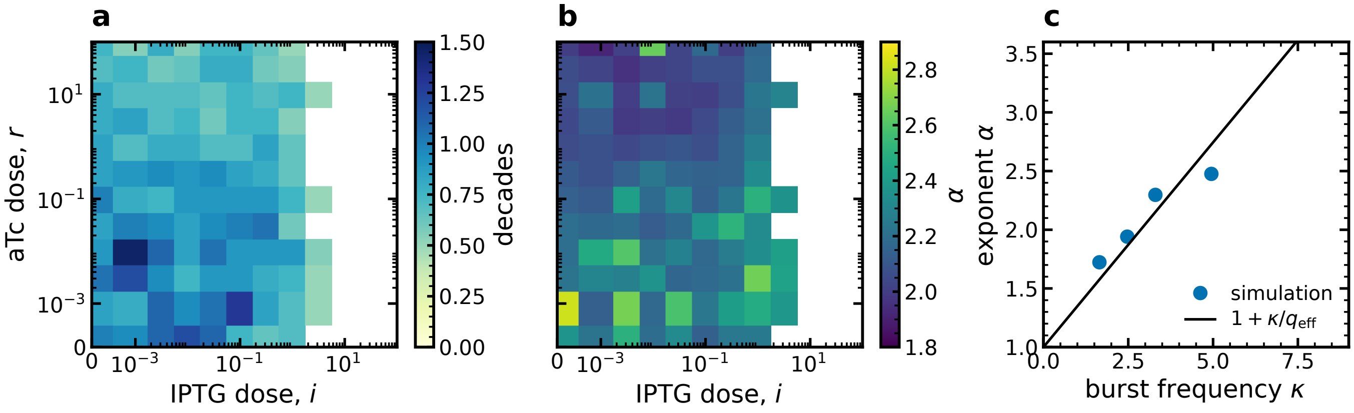


FIG. S2. **Tails over the** IPTG-aTc **plane and the role of the sensor.** (a) Scaling range of the reporter distribution in decades over the plane of IPTG and aTc doses; white, no scaling range of at least half a decade. (b) Exponent of the scaling range where it exists; each cell is one exact simulation of the closed loop on the dose grid. (c) Exponent against the burst frequency $\kappa$ of the *lacI* gene, with the prediction of Eq. (17) (one fitted constant $q_{\mathrm{eff}}$); larger $\kappa$ has no resolvable scaling range. Exact simulations of the closed loop.

A gene that fires in bursts of mean size $b_I$ has an exponent set by its firing frequency $\kappa$, not by its message rate $b_I\kappa$. At the same firing frequency, bursts of three messages give an exponent of 2.47 against 2.32 for single messages, and at the same message rate the exponent falls to 1.60, in line with Eq. (17) (Table S3).

Two controls isolate the loop. A reporter without repression, at the same deterministic demand, has no scale-free range: its normalized fluorescence has a coefficient of variation of 0.07, and its traffic intensity moves in proportion to the drive. A *lacI* message that lacks the target is blind to the queue; its protein sits at $\Lambda$ and silences the reporter, whose median message count is 0; rare escapes give its fluorescence a mean-to-median ratio of 11. This control has a broad scaling range too (2.5 decades at exponent 1.8), but around a silent reporter: the scale-free excursions are the sensor's, and reading the queue sets the baseline at the critical point. The queue also enters the exponent through the bracketed factor of Eq. (16): in the closed loop the reporter carries nearly all of the demand, so the factor runs from one in congested excursions to its maximum at the set point, whereas the silent reporter of the blind-sensor control carries almost none and the factor is close to one.

| Variant | $\rho_{\mathrm{det}}$ | $\alpha$ (EGFP) | range (decades) | 99th pct. / median |
|---|---|---|---|---|
| reference | 0.95 | 2.31 | 1.03 | 27 |
| $n_I = 1.5$ ($\Lambda = 125$ at fixed $\rho_{\mathrm{det}}$) | 0.95 | 2.55 | 0.77 | 17 |
| $n_I = 3$ ($\Lambda = 26$ at fixed $\rho_{\mathrm{det}}$) | 0.95 | 1.94 | 1.05 | 54 |
| bursty reporter gene (mean burst 5) | 0.95 | 2.31 | 1.00 | 27 |
| slower messages ($\varepsilon = 0.2$) | 0.95 | 2.23 | 0.82 | 26 |
| faster messages ($\varepsilon = 0.05$) | 0.95 | 2.46 | 0.87 | 27 |
| $\Omega = 3$ | 0.95 | 2.19 | 1.25 | 34 |
| $\Omega = 30$ | 0.95 | 2.22 | 0.92 | 27 |
| weaker promoter ($\alpha_G = 650$) | 0.69 | 2.00 | 0.85 | 34 |
| stronger promoter ($\alpha_G = 1.04 \times 10^4$) | 1.28 | 2.38 | 1.12 | 22 |
| LacI gene, single messages ($\kappa = 3.3$) | 0.95 | 2.32 | 1.10 | 26 |
| LacI gene in bursts of 3, same event rate ($\kappa = 3.3$) | 0.95 | 2.47 | 1.05 | 27 |
| LacI gene in bursts of 3, same message rate ($\kappa = 1.1$) | 0.95 | 1.60 | 1.18 | 104 |

TABLE S3. **Sensitivity of the exponent.** Closed loop at zero inducers, EGFP filter; mean of three runs of 12 000 dilution lifetimes each (4000 for $\Omega = 30$), with the exponent averaged over the runs that have a scaling range and the range over all runs. Other cooperativities are given the $\Lambda$ that keeps the reference $\rho_{\mathrm{det}}$. The last three rows are means of two runs of 8000 lifetimes of a *lacI* gene that fires in single messages or in bursts of three; $\kappa$ is the frequency of firing.

Across the model variants in Table S3, the exponent changes by a few tenths and the scaling range remains about a decade. The variants change cooperativity at the same deterministic operating point, message lifetime, counting scale or promoter strength.

#### S4. PREDICTIONS THAT SEPARATE SELF-ORGANIZATION FROM REPRESSION

The distributions above are properties of a state; self-organization is a property of a response, and two perturbations expose it (Fig. S3). A reporter under a LacI-like repressor that does not read the queue falls when the silencing capacity rises. In the loop the opposite happens: more capacity lowers the LacI message lifetime, releases the promoter, and demand rises to refill the enlarged queue, so that $\rho$ stays near one and the reporter follows the capacity. At fixed $\Gamma$ the reporter message is

$$G^* = A_G\, W \simeq \frac{\rho\sqrt{\eta}}{\Gamma}, \tag{S17}$$

so for $n_I = 2$, where $\Gamma$ does not depend on the capacity, the median reporter follows $\sqrt{\eta}$ [Fig. S3(a)]. From zero to saturating aTc it should rise by the factor

$$\sqrt{\frac{\eta_\infty}{\eta_0}} = 2.0, \tag{S18}$$

and the simulated median rises by 2.2. After a jump in aTc the reporter first dips, because message clearance responds before the sensor does, then recovers and settles above its initial level, whereas without the loop it falls monotonically [Fig. S3(d)]. In IPTG the loop buffers the response while $\Gamma$ is in the band [Fig. S3(b)].

The second perturbation is a step of IPTG, the one inducer that acts on demand rather than on capacity. By Eq. (13), IPTG enters the operating point only through $1 + i$. Let

$$f = \frac{1 + i'}{1 + i} \tag{S19}$$

be the factor by which a change of the dose from $i$ to $i'$ multiplies $1 + i$. It multiplies the drive of a repressed reporter promoter by $f^{n_I}$, the same as a change of the promoter strength by that factor at fixed controller, without changing the construct. A reporter promoter whose repressor does not read the queue follows it, so the traffic intensity of the blind sensor rises by the factor $f^{n_I}$ and stays there. With the loop, $\Gamma$ is only divided by $f$, so the median rises at once and is then compressed within a fraction of a dilution lifetime to

$$\rho - 1 \simeq \frac{f\,\Gamma_0^{-1} - \Gamma_0\, f^{-1}}{\sqrt{\eta}} \tag{S20}$$

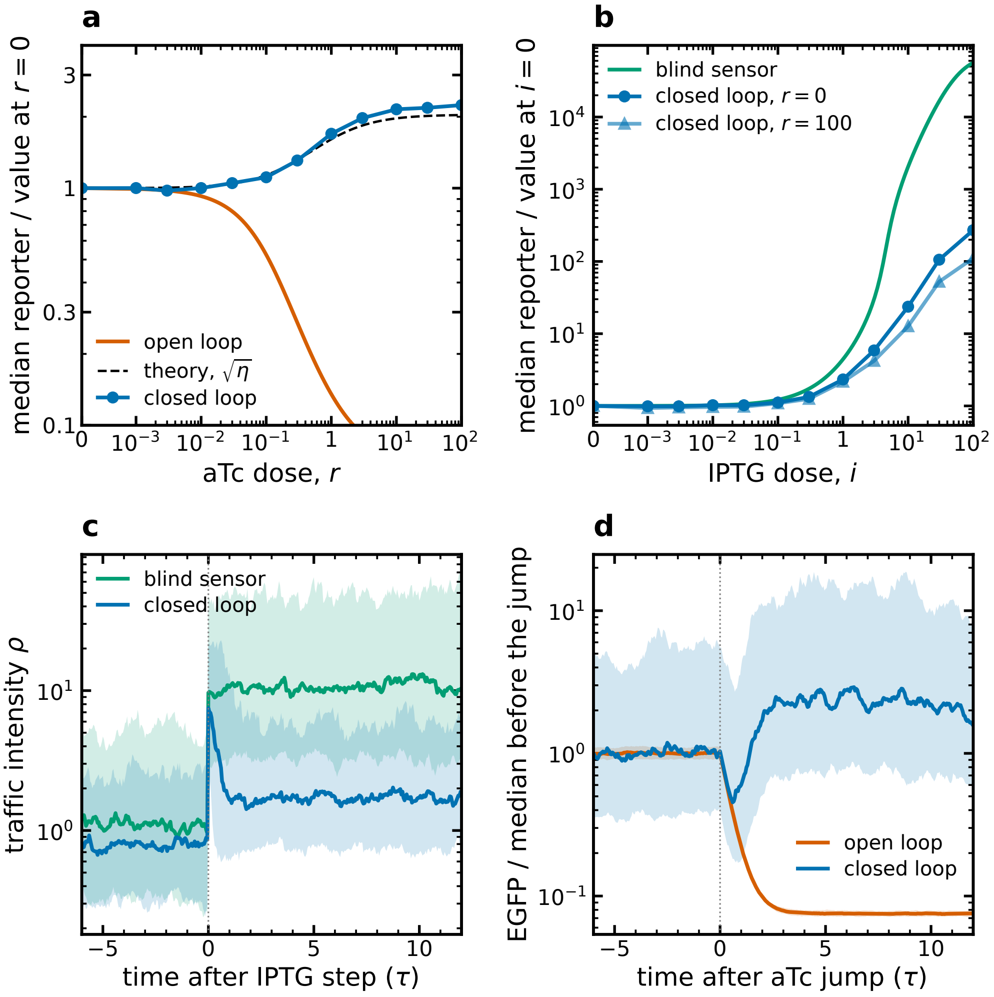


FIG. S3. **Predicted responses that a repressor alone cannot give.** (a) Median reporter against aTc dose at zero IPTG, relative to its value at zero aTc: with the loop it rises as the square root of the capacity (dashed line, theory), without it it falls. (b) Median reporter against IPTG dose, relative to its value at zero IPTG, at two aTc doses, and for a sensor blind to the queue. (c) Traffic intensity after a step of IPTG from zero to a dose that multiplies the reporter drive tenfold, at the time marked by the dotted line (median and 10-90 % range of the cells), for the closed loop and for a sensor blind to the queue whose reporter drive is set to the same initial traffic intensity. (d) EGFP reporter after a jump of aTc from zero to saturation, normalized by its median before the jump. Time $\tau$ is in dilution lifetimes.

[Fig. S3(c)]. The open loop, whose promoter is not repressed, ignores IPTG and is the control for the aTc jump instead. Both perturbations use only the inducers of the design and the same lines and detector as Fig. 5. The step is limited by how fast IPTG reaches LacI, minutes to 1 h, which is short compared with a dilution lifetime.

#### S5. DATA, TAIL STATISTICS AND REPORTER CALIBRATION

*a. Imaging.* Cells of both lines were also imaged by holotomography on a Nanolive microscope, at the EMBL Mesoscopic Imaging Facility. Holotomography is a label-free, quantitative-phase technique: the instrument records holograms of the sample under many illumination angles and reconstructs from them the three-dimensional refractive index (RI) of the cell, which reports its dry mass density without staining. The RI channel in Figs. 4 and S4 is the maximum-intensity projection of this volume along the optical axis (RI MIP), which outlines the cell body, the nucleus and its nucleoli. GFP was recorded with the fluorescence module of the same instrument (FITC channel), so that fluorescence and RI are registered in the same field. For the overlay in Fig. 4, the smooth background of each fluorescence image was estimated by a morphological opening and subtracted, the remaining noise floor (the median plus three robust standard deviations of the background pixels) was removed. A single linear intensity scale, set by

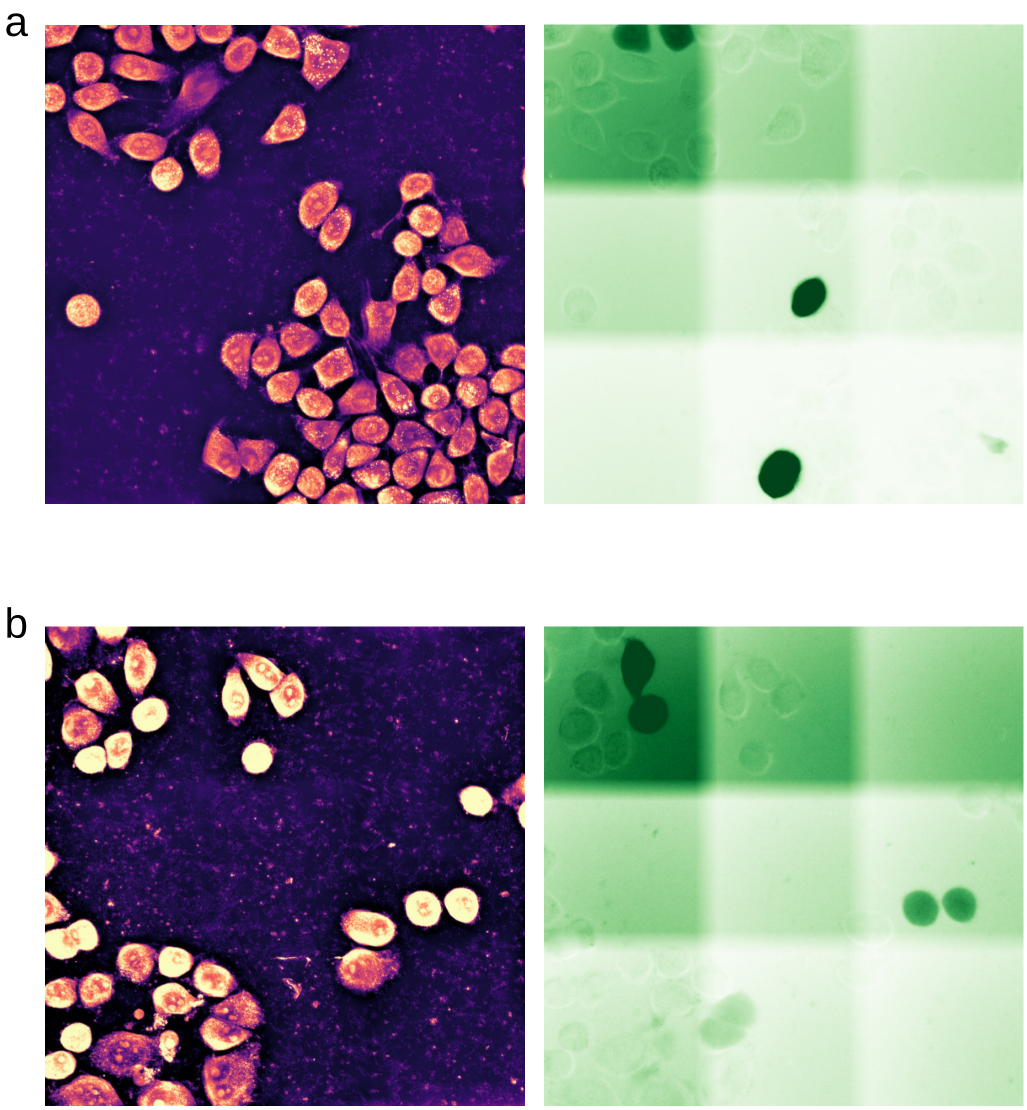


FIG. S4. **Unmerged channels of Fig. 4.** Refractive index (RI MIP, left; lighter is higher) and GFP fluorescence (FITC, right; darker green is brighter) for the (a) d2EGFP and (b) EGFP lines. The FITC images are shown with their raw intensity scaling.

the brightest remaining pixel over the fields shown, was applied to both lines. The fluorescence images were used only to show that reporter intensities differ between cells; quantitative distributions come from flow cytometry (Fig. 5).

*b. Conventional tail fits.* Fitting a Pareto tail above the Kolmogorov-Smirnov-optimal threshold gives exponents of 2.68 and 2.19 for EGFP and d2EGFP. These estimates use millions of cells, but cell-level precision does not represent clone-to-clone uncertainty; their spread is also sensitive to threshold choice (removing the top 1 % of events gives 2.90 and 2.68). We therefore report the exponent of the scaling range, the quantity that the same procedure gives for the model. Both tails are better described by a power law with an exponential taper (cutoffs at 55 and 152 times the threshold) than by an untruncated power law ($\Delta$AIC $= 11$ and 209), a lognormal (5 and 116; the EGFP margin is small because its threshold leaves only 0.5 % of the events) or an exponential. On a seeded subsample of at most 5000 events, a 99-replicate threshold-refitted Pareto goodness-of-fit bootstrap gives $p = 0.01$ for EGFP and $p = 0.07$ for d2EGFP. At a 0.05 threshold this rejects the untruncated Pareto for EGFP but not for d2EGFP; the p-value resolution is 0.01. The Pareto exponents of the two halves of each acquisition differ by at most 0.03.

*c. Reporter calibration.* For physical time, the loss rate in units of the dilution rate is

$$\delta_F = \frac{k_{\text{eff}}}{\delta_c}, \qquad k_{\text{eff}} = k_{\text{deg}} + \delta_c. \tag{S21}$$

After protein production has stopped and maturation is complete, a decay experiment gives

$$F(t) = F_{\mathrm{bg}} + F_0 e^{-k_{\mathrm{eff}} t}, \qquad k_{\mathrm{deg}} = k_{\mathrm{eff}} - \frac{\ln 2}{T_d}, \tag{S22}$$

with the doubling time $T_d$ measured under the assay conditions [1]. Stopping transcription alone leaves translation from residual message, so the fitting window must lie after it has decayed. An assay-specific decay curve can be fit with this relation to estimate $\delta_F$ and separate protein degradation from dilution.

[1] P. Corish and C. Tyler-Smith, Protein Engineering, Design and Selection **12**, 1035 (1999).
[2] X. Li, X. Zhao, Y. Fang, X. Jiang, T. Duong, C. Fan, C.-C. Huang, and S. R. Kain, Journal of Biological Chemistry **273**, 34970 (1998).
